# Surface-driven RNA-refolding by the OB-fold proteins of the *Trypanosoma brucei* editosome

**DOI:** 10.1101/099705

**Authors:** Christin Voigt, Mateusz Dobrychłop, Elisabeth Kruse, Anna Czerwoniec, Joanna M. Kasprzak, Patrycja Bytner, Janusz M. Bujnicki, H. Ulrich Göringer

## Abstract

RNA editing in African trypanosomes represents an RNA-processing reaction that generates functional mitochondrial transcripts from sequence-deficient pre-mRNAs. The reaction is catalyzed by a macromolecular protein complex known as the editosome. Editosomes have been demonstrated to execute RNA-chaperone activity to overcome the highly folded nature of pre-edited substrate mRNAs. The molecular basis of this activity is unknown. Here we test five OB-fold proteins of the editosome as potential candidates. We show that the different proteins interact by hetero-oligomerization and we demonstrate that all proteins execute RNA-chaperone activity. Activity differences correlate with the surface areas of the proteins and map predominantly to the intrinsically disordered subdomains of the polypeptides. To provide a structural context for our findings we present a coarse-grained model of the editosome. The model suggests that an inner core of catalytically active editosome components is separated from an outer shell of IDP-domains that act as RNA-remodeling sites.

## INTRODUCTION

Editosomes are high molecular mass (0.8MDa, 20S) protein complexes that catalyze the U-insertion/deletion RNA-editing reaction in African trypanosomes and other kinetoplastid organisms^1,2^. The processing reaction is characterized by the site-specific insertion and to a lesser extent, deletion of exclusively U-nucleotides and converts cryptic primary transcripts into translatable mRNAs. Editosomes harbor one substrate-RNA binding site^3^ and interact with a large set of pre- and partially edited mitochondrial transcripts to execute the reaction. Recent structure probing experiments uncovered that the pre-edited mRNAs are unusually folded. The different transcripts have thermodynamic stabilities that resemble structural RNAs^4^ and due to a very high content of runs of G-nucleotides they contain multiple, up to five, G-quadruplex (GQ)-folds^5^. Perhaps as a consequence, *T. brucei* editosomes execute a complex-intrinsic RNA-chaperone activity^3^. The activity acts by increasing the flexibility of predominantly U-residues to lower their base-pairing probability thereby generating a simplified RNA-folding landscape with a reduced energy barrier to facilitate the binding of gRNAs^4^. Thus, the editosome-driven RNA-unfolding reaction is important for the editing cycle especially during the initiation and elongation phases of the process. However, the molecular nature of the chaperone activity is not understood.

Proteins with RNA-chaperone activity represent a structurally diverse group of polypeptides that contribute to almost all RNA-driven biochemical processes in all domains of life^6^. This includes many viral and bacterial proteins such as Ncp7^7^, StpA and Hfq^8,9^ and a large number of ribosomal proteins of both, pro- and eukaryotic origin^10^. Importantly, several of the RNA chaperones contain one or more OB-fold motif(s). OB refers to the general oligonucleotide/oligosaccharide-binding ability of the proteins, which is mediated by a five-stranded β-barrel fold^11^. OB-folds have specifically been identified in bacterial and plant cold shock proteins (Csp) where they contribute to resolve misfolded RNA-species^12,13^. Furthermore, OB-fold proteins are universally involved in RNA-remodeling steps prior to the initiation of protein biosynthesis with ribosomal protein S1 and *E. coli* initiation factor 1 (IF1) containing multiple OB-domains^14,15^.

The protein inventory of the *Trypanosoma brucei* 20S editosome lists six OB-fold proteins termed TbMP81, TbMP63, TbMP42, TbMP24, TbMP19 and TbMP18 following the nomenclature of Worthey et al., 2003^16^. TbMP stands for *Trypanosoma brucei* mitochondrial protein followed by a number indicating the calculated molecular mass in kDa. TbMP81, TbMP63 and TbMP42 additionally contain two C2H2-type Zn-fingers or C2H2-Zn-finger-like domains and all six proteins have recently been predicted to harbor long stretches of intrinsically disordered regions (IDR)^17^. The domain structure of the different proteins, a 3D-consensus model of the OB-fold and a plot of their disorder propensities^18^ is summarized in Fig. 1. While the percentage of predicted disorder varies between 25% in TbMP18 and 72% in TbMP81, together, the six proteins are more disordered than the average of all other editosomal proteins^17^. Two of the proteins (TbMP63, TbMP81) even contain more disordered than ordered regions.

**Figure 1.**
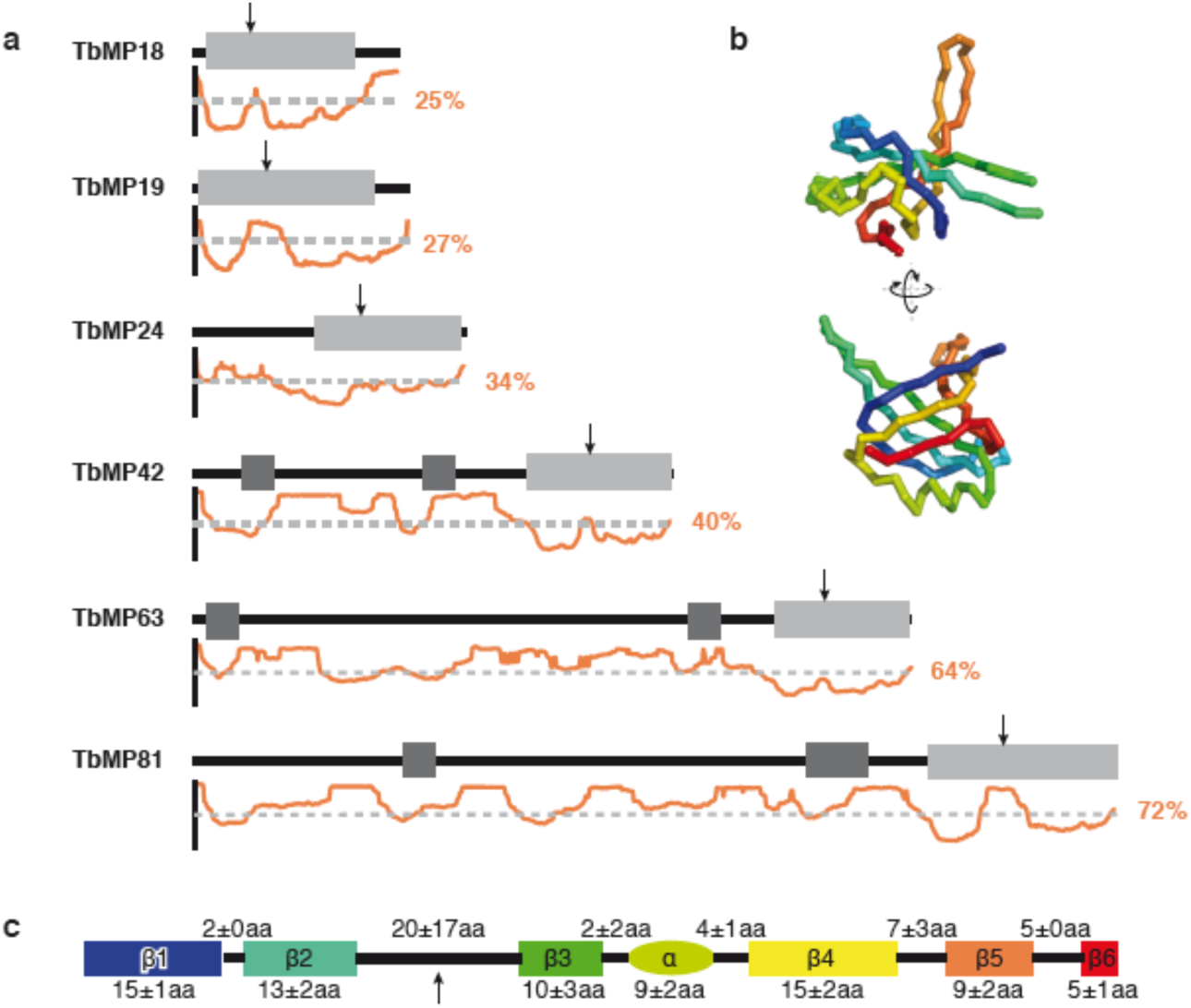
Domain structure of the OB-fold proteins of the *T. brucei* editosome. (a) OB-fold domains: boxes in light grey. C2H2-zinc fingers and C2H2-zinc finger-like domains: boxes in dark gray. Arrow: Position of the loop (L)-sequence between β-sheets β2 and β3 (L_23_). Disorder propensity plots (between 0 and 1) for each protein are shown in orange. The grey dashed line indicates a value of 0.5. Average disorder propensities are listed as percent values. (b) Consensus structure of all editosomal OB-folds in spectral colors from the N-terminus in blue to the C-terminus in red. For details see Supplementary Table S1. (c) Sketch of the secondary structural elements (colored as in B) of the consensus OB-fold structure. Rectangles: β-sheets. Oval sphere: α-helix. Black lines: loop regions. Numbers: average amino acid length (±SD). Details are given in Supplementary Table S2.

Although several of the editosomal OB-fold proteins have been investigated in gene ablation experiments and as recombinant proteins, no coherent function for any of the six polypeptides has emerged from these studies. TbMP24, TbMP18 and TbMP42 were identified as RNA-binding proteins^19–21^. Recombinant TbMP42 was characterized to execute endo/exonuclease activity^21,22^ and recombinant TbMP24 was shown to perform RNA-annealing activity^23^. Gene knockdown of five of the proteins (TbMP18, TbMP24, TbMP42, TbMP63, TbMP81) identified an impact on the structural integrity of the editosome^20,24–27^, which was supported by yeast two-hybrid experiments in combination with protein/protein interaction studies^28^. The data suggest a scenario in which TbMP18, TbMP24, TbMP42, TbMP63 and TbMP81 interact with each other, mainly relying on their OB-fold domains thereby forming a clustered “OB-fold core” within the 20S editosome^29^. Recent interprotein chemical cross-linking data support the existence of such a “core-domain” and have identified cross-links between all six OB-fold proteins most of which within or in proximity to the OB-fold domains^30^.

Importantly, clustered OB-folds in the yeast RRP44 protein have been demonstrated to catalyze the unwinding of dsRNA^31,32^. As a consequence, Böhm et al., 2012^3^ suggested that the potential OB-fold core of the *T. brucei* editosome might act in a similar fashion. Here we show that the *T. brucei* editosomal OB-fold proteins indeed catalyze a RNA-refolding reaction. We demonstrate that the activity is inherent to the intrinsically disordered protein (IDP)-regions of the different proteins and we uncovered a correlation of the chaperone activity to the surface areas of the proteins. Using a coarse-grained modeling approach we provide a molecular model of the *T. brucei* 20S editosome, which suggests that the high molecular mass complex has a bi-partite composition: an outer shell of intrinsically disordered proteins that act as RNA-contact and RNA-remodeling elements and an inner core of structurally defined proteins, mediating the catalytic reactions of the complex.

## RESULTS

### Recombinant expression of the OB-fold proteins of the *T. brucei* editosome

To test whether the editosome-inherent RNA-chaperone activity is mediated by one or several of the editosomal OB-fold proteins we expressed the six proteins as recombinant polypeptides in *E. coli*. TbMP81, TbMP63, TbMP42 and TbMP24 were expressed as full-length (FL) and as OB-fold-only (OB) constructs. TbMP19 and TbMP18, due to their small size, were expressed as FL-constructs only. While the different constructs expressed well upon induction, the majority of polypeptides remained insoluble at an expression temperature of 37°C. Lowering the temperature to 18°C improved the solubility, however, all attempts to express soluble TbMP19-FL and TbMP63-FL failed. For TbMP63-OB and TbMP24-FL we devised a co-expression regime together with TbMP18-FL as a previously identified interaction partner^28^. In the end 8 of the 10 recombinant protein constructs were available in yields between 5-40mg soluble protein/L *E. coli* culture (Fig. 2a and Supplementary Table S4).

**Figure 2.**
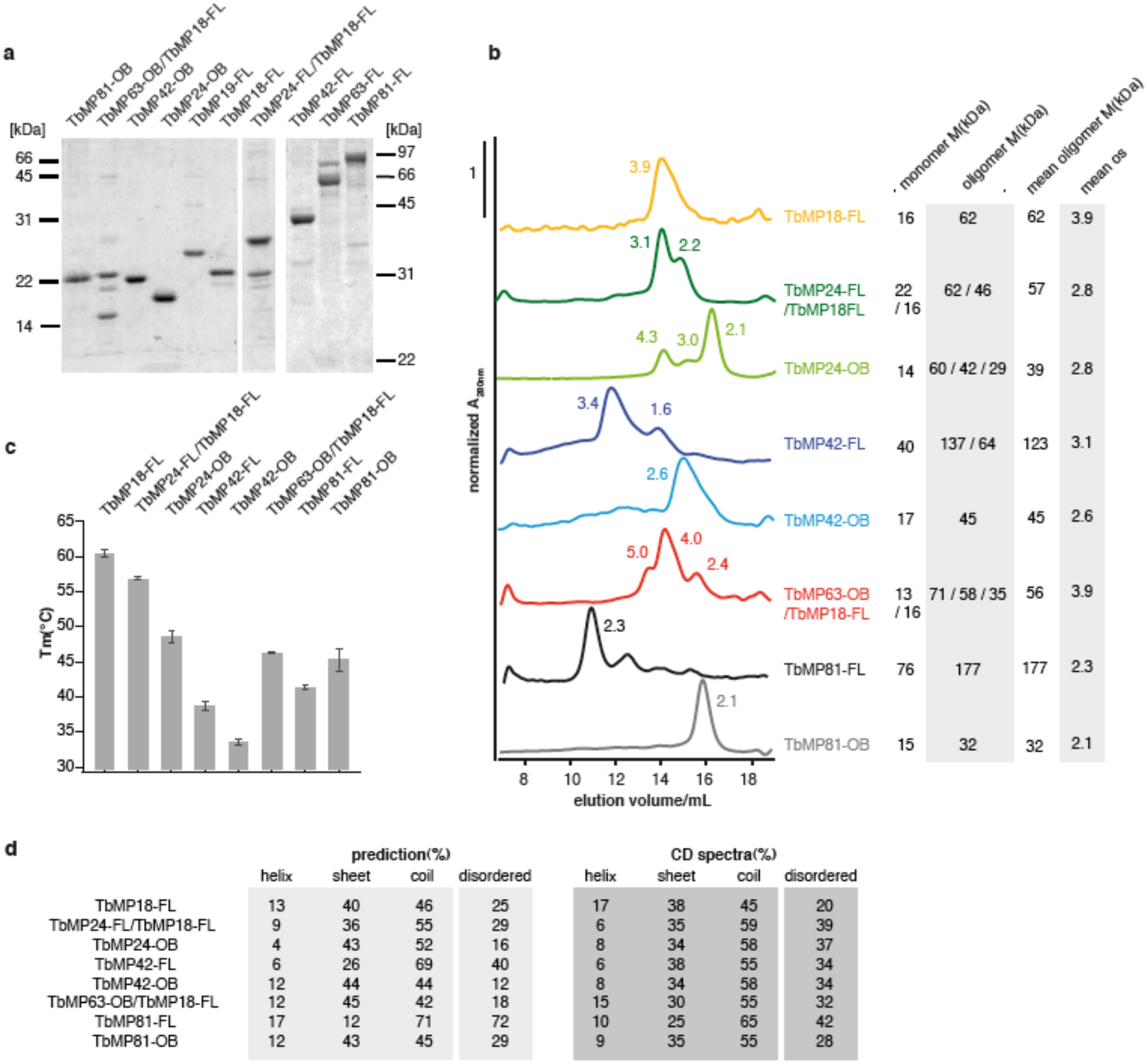
Characterization of recombinant editosomal OB-fold proteins. (a) Gel electrophoretic analysis of purified protein preparations in 15% (w/v) (left) and 10% (w/v) (right) SDS-containing polyacrylamide gels. Apparent molecular masses are in kDa. (b) Normalized size exclusion chromatography profiles of purified OB-fold proteins. Peak numbers indicate oligomerization state(s) (os), which were determined as outlined in the Experimental Procedures section. The molar ratios of the coexpressed protein constructs were determined as 2:1 and 1.5:1 for the TbMP24-FL/TbMP18-FL construct and as 3:2, 2:2 and 1:1 for the TbMP63-OB/TbMP18-FL complex. (c) Differential scanning fluorimetry (DSF)-based melting transitions (T_m_) of the different protein constructs. Error bars are SD. For an example see Supplementary Fig. S1. (d) Circular dichroism (CD)-derived secondary structure contents of recombinant OB-fold proteins. Predicted secondary structure contents were derived from the homology models (see Supplementary Tables S1 and S2). The structure of non-modeled protein domains was predicted using PSIPRED^55^. The molar ratios of the co-expressed constructs was determined as 1.1:1 for the TbMP24-FL/TbMP18-FL complex and as 1:1.5 for the TbMP63-OB/TbMP18-FL complex (see also Supplementary Table S4). Disorder propensities were calculated using MetaDisorderMD2^18^.

Notably, all expressed protein constructs formed multiple homo-oligomeric higher order assemblies with one dominant assembly state (Fig. 2b). TbMP81-FL and TbMP81-OB predominantly formed homodimers, TbMP42-FL, TbMP42-OB and TbMP24-OB homotrimers and TbMP18-FL homotetramers. Importantly, the identified oligomers of the FL and OB-only constructs were invariably identical, indicating that the interaction surfaces involve the OB-fold domains of the different proteins. In the case of the two co-expressed constructs (TbMP24-FL/TbMP18-FL, TbMP63-OB/TbMP18-FL) we exclusively identified hetero-oligomeric assemblies (Fig. 2b), suggesting an enhanced stability and/or solubility of the hetero-oligomeric complexes over the homo-oligomeric complexes.

### Structural characteristics of the recombinant OB-fold proteins

The purified protein preparations were analyzed for their secondary (2D)-structure content using circular dichroism (CD) spectroscopy (Fig. 2d). As expected and in line with structure prediction algorithms, all proteins display a high fraction of unstructured sequence stretches. The α-helical content of the different proteins is as low as 6-17%, the β-sheet content varies between 25-38% and the amount of coil structures varies between 45-65%. On average 33% of the protein sequences are disordered. To further assess the characteristics of the proteins, we measured their thermal unfolding using differential scanning fluorimetry^33^. Intrinsically disordered proteins typically display broad denaturation profiles and low cooperativity in the folding/unfolding transition^34^. All melting curves were converted into fraction folded (α) *vs*. temperature plots to derive half-maximal melting temperatures (T_m_) (Supplementary Fig. S1) and the data are summarized in Fig. 2c. Despite the fact that all proteins contain a similarly structured OB-fold, they cover a broad thermal stability range: The OB-fold-only construct of TbMP42 is the least stable polypeptide (T_m_ 34°C) and TbMP18-FL represents the most stable protein (T_m_ 61°C). Furthermore, by comparing the FL and OB-fold-only versions of the same proteins we uncovered that OB-folds can be stabilized as well as destabilized by the remaining amino acid sequences. TbMP42-OB is stabilized with a ΔT_m_ of 5.2°C and TbMP81-OB is destabilized with a ΔT_m_ of 3.9°C. Thus, sequences outside the OB-folds modulate the thermal stability of the different protein constructs. The same holds true for the bimolecular complexes TbMP18-FL/TbMP24-FL and TbMP18-FL/TbMP63-OB. Both complexes have a lower T_m_ than TbMP18-FL alone.

### The formation of hetero-oligomeric complexes

Editosome subcomplex reconstitution experiments as well as recent chemical cross-linking experiments demonstrated that the OB-fold proteins of the *T. brucei* editosome interact with each other^28–30^. This tempted us to analyze the association behavior of the different protein preparations in “mix-and match-type” binding experiments. Fig. 3a shows as a representative example the formation of a trimeric (1:2) TbMP18-FL/TbMP24-OB complex. Pairwise interaction experiments were conducted with all combinations of proteins and we identified hetero-oligomers for TbMP18-FL with TbMP24-OB, TbMP42-FL, TbMP42-OB and TbMP81-FL (Fig. 3b, d). Importantly, no pairwise interaction was detected without TbMP18-FL. Furthermore, TbMP42-FL as well as TbMP42-OB were able to hetero-oligomerize with TbMP18-FL, again indicating that the TbMP42-OB-fold is sufficient to mediate the interaction. The same holds true for the association of TbMP24 with TbMP18. Thus, the individual OB-folds likely act as docking modules.

**Figure 3.**
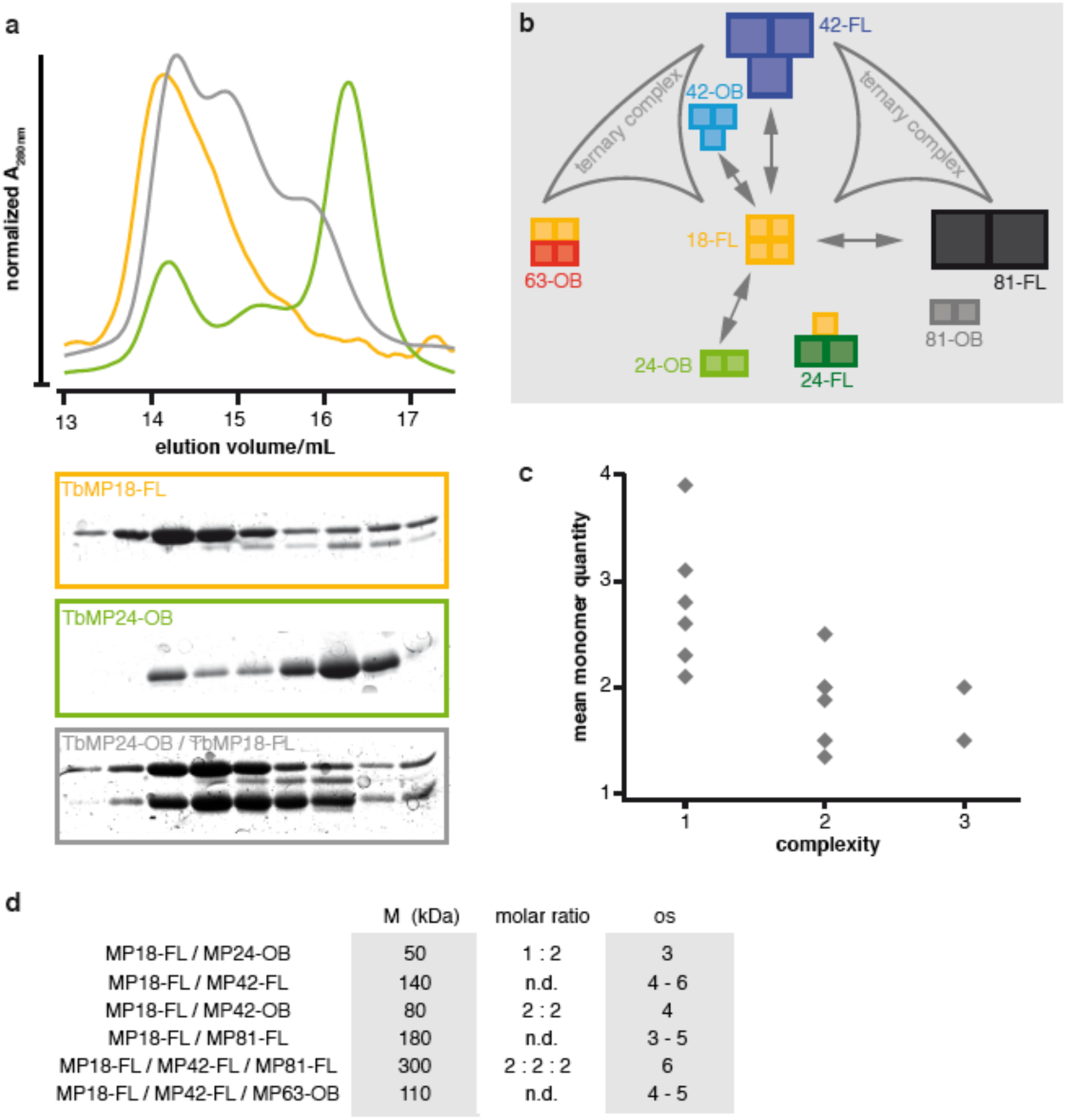
The formation of OB-fold protein complexes *in vitro*. (a) Normalized size exclusion chromatography profiles(upper panel) and gel electrophoretic analysis (lower panels) of a representative experiment to demonstrate the formation of the hetero-oligomeric TbMP24-OB/TbMP18-FL complex (grey). Green: TbMP24-OB. Yellow: TbMP18-FL. See also Supplementary Fig. S2. (b) *In vitro* interaction map. Summary of all identified protein interactions (arrows). The size of the colored boxes indicates the apparent molecular sizes of the different proteins and the number of boxes refers to the number of monomeric units in the most abundant complex. Curved triangles: ternary interactions. (c) Correlation of the mean number of protein monomers in all homo- and hetero-oligomeric complexes as a function of the complexity of the complexes with respect to the number of unique subunits. (d) Summary of the characteristics of the hetero-oligomeric complexes. os: oligomerization state. n.d.: not determined. See also Supplementary Fig. S2.

We also identified two ternary complexes: First, a complex between TbMP42-FL, TbMP18-FL and TbMP63-OB, which was identified before^28^ and second, a ternary assembly between TbMP81-FL, TbMP42-FL and TbMP18-FL (Supplementary Fig. S2). The two ternary complexes are mutually exclusive since we were not able to identify a complex containing both, TbMP81-FL and TbMP63-OB. Also, no ternary complex containing TbMP24 was found. Importantly, all heteromeric complexes contain less identical monomer subunits when compared to the corresponding homomeric complexes (Fig. 3c). Thus, the proteins interact by re-organizing the single monomers instead of adding up the different homo-oligomers to larger heteromeric assemblies. A summary of the data is shown in Fig. 3b,d.

### The OB-fold proteins of the *T. brucei* editosome execute RNA-remodeling activity

To quantitatively assess the potential RNA-remodeling activity of the different OB-fold proteins we relied on the recently described RNaseH-based guide (g)DNA-annealing assay^4^. The assay is based on the rationale that a remodeled *i.e.* structurally open target RNA should bind a guide RNA-mimicking, short complementary DNA-oligonucleotide more readily when compared to a structurally constrained RNA. The formed pre-mRNA/DNA-oligonucleotide hybrid molecules were identified by RNaseH cleavage followed by an electrophoretic separation of the resulting RNA-fragments (Fig. 4a). As a representative *T. brucei* mitochondrial transcript we used the pre-edited mRNA of apocytochrome b (CYb). The RNA-chaperone activity of 20S editosomes served as a positive control^3,4^. Importantly, the experiments were performed with varying amounts of protein to use the protein concentration required to achieve half-maximal cleavage (c_1/2_) as a metric for the RNA-remodeling activity.

**Figure 4.**
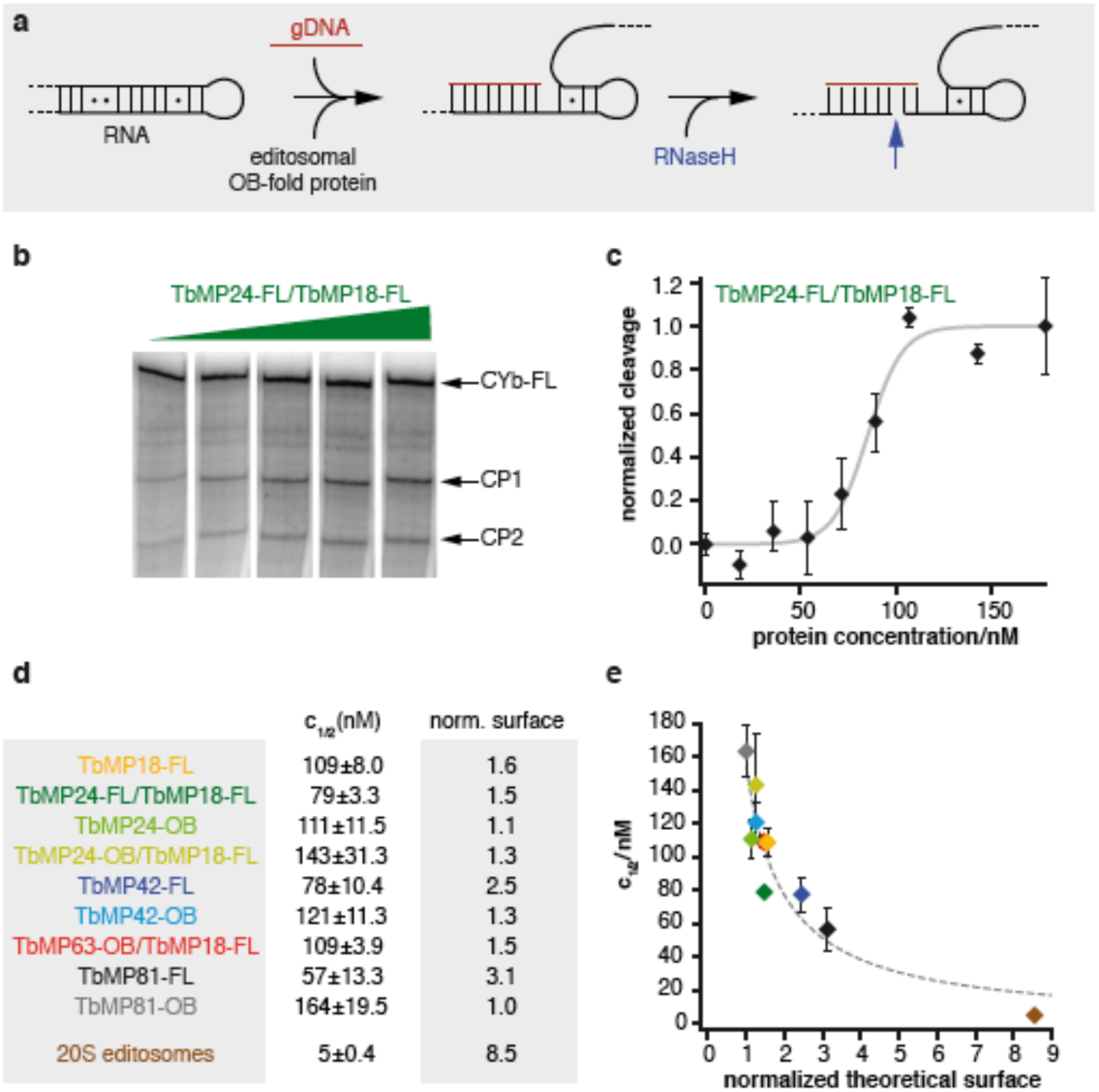
RNA-chaperone activity of the editosomal OB-fold proteins. (a) Sketch of the RNaseH-based “gDNA”-annealing assay^4^. (b) Representative gel electrophoretic separation of RNaseH-induced RNA-cleavage products (CP1/CP2) of pre-edited CYb-mRNA using increasing concentrations of TbMP24-FL/TbMP18-FL: 0, 70, 110, 140, 210nM (left to right). Cyb-FL: full length CYb-pre-mRNA. (c) Representative plot of normalized cleavage values as a function of the TbMP24-FL/TbMP18-FL concentration. Grey line: Sigmoidal fit of the data to derive half-maximal cleavage concentrations (c_1/2_) as a metric for the RNA-chaperone activity. (d) Summary of all c_1/2_-values (±SD) and normalized surface areas of the tested editosomal OB-fold constructs in comparison to 20S editosomes. (e) Plot of the derived c_1/2_-values of all tested OB-fold complexes in relation to their normalized surface areas. Dotted line in grey: Reciprocal fit of the data. Error bars are SD-values.

All purified OB-fold homo-oligomers, co-expressed hetero-oligomers as well as the *in vitro* formed TbMP24-OB/TbMP18-FL hetero-oligomer were tested. Fig. 4b, c show a representative example for the TbMP24-FL/TbMP18-FL complex. All data are summarized in Fig. 4d. Surprisingly, all protein constructs show RNA-remodeling activity. The measured c_1/2_-values range from 57±13nM (TbMP81-FL) to 164±16nM (TbMP81-OB) with a mean of 108±12nM. This represents a roughly 20-fold higher protein concentration in comparison to 20S editosomes (c_1/2_ 5±0.4nM). Thus, although all protein constructs are capable of remodeling the pre-mRNA, they are less active than 20S editosomes. This suggests that the RNA-remodeling activity of the editosome likely represents a cumulative trait, to which each of the different OB-fold proteins contributes a defined increment. Furthermore, we noticed that all FL-proteins are more active than the OB-fold-only constructs. This indicates that protein sequences outside the OB-fold domains, *i.e.* the intrinsically disordered regions of the proteins are for the most part responsible for the remodeling activity.

Importantly, for intrinsically disordered RNA-binding proteins it was shown that the size of the RNA-binding surfaces correlate with the overall sizes of the proteins ^35^. As a consequence, we analyzed whether the RNA-remodeling activities of the different OB-fold proteins can be correlated to their surface areas. For that we calculated relative surface areas for all proteins and plotted them in relation to the measured RNA-remodeling activities (c_1/2_-values). Fig. 4e shows the resulting plot. The two variables show a reciprocal dependency and the data fit to a hyperbole with a correlation coefficient (*r^2^*) of 0.87. Thus, a major determinant of the RNA-chaperone activity of the different OB-fold proteins is their surface area. Proteins with larger surfaces such as TbMP81-FL execute a higher refolding activity and *vice versa*. The high remodeling activity of the editosome (c_1/2_ 5±0.4nM) is the result of the large surface of the 0.8MDa complex.

### Structural characterization of the RNA-remodeling activity

As a follow up of the results above we asked the question how the RNA-remodeling activities of the different OB-fold proteins manifest on a structural level and how they compare to the structural changes induced by 20S editosomes^4^. For that, we mapped the chaperone-induced structure changes with nucleotide resolution using SHAPE (Selective 2’-Hydroxyl Acylation analyzed by Primer Extension) chemical probing^36,37^. TbMP18-FL was selected as a representative OB-fold protein and the pre-edited mRNA of RPS12 as a typical *T. brucei* mitochondrial transcript. Fig. 5a shows the SHAPE-reactivity profile of the RNA in the presence of TbMP18-FL. As anticipated, the protein induces an 18% increase in the mean SHAPE-reactivity (Fig. 5b), which is in line with a remodeling reaction similar to 20S editosomes (26%). Twenty two percent of the nucleotide positions in the RPS12-transcript are responsive to the refolding reaction and as evidenced in the difference (Δ) SHAPE-profile (Fig. 5c, d), the majority of affected nucleotides (71%) increase in flexibility. Again, this represents a value comparable to the remodeling activity of the 20S editosome (72%).

**Figure 5.**
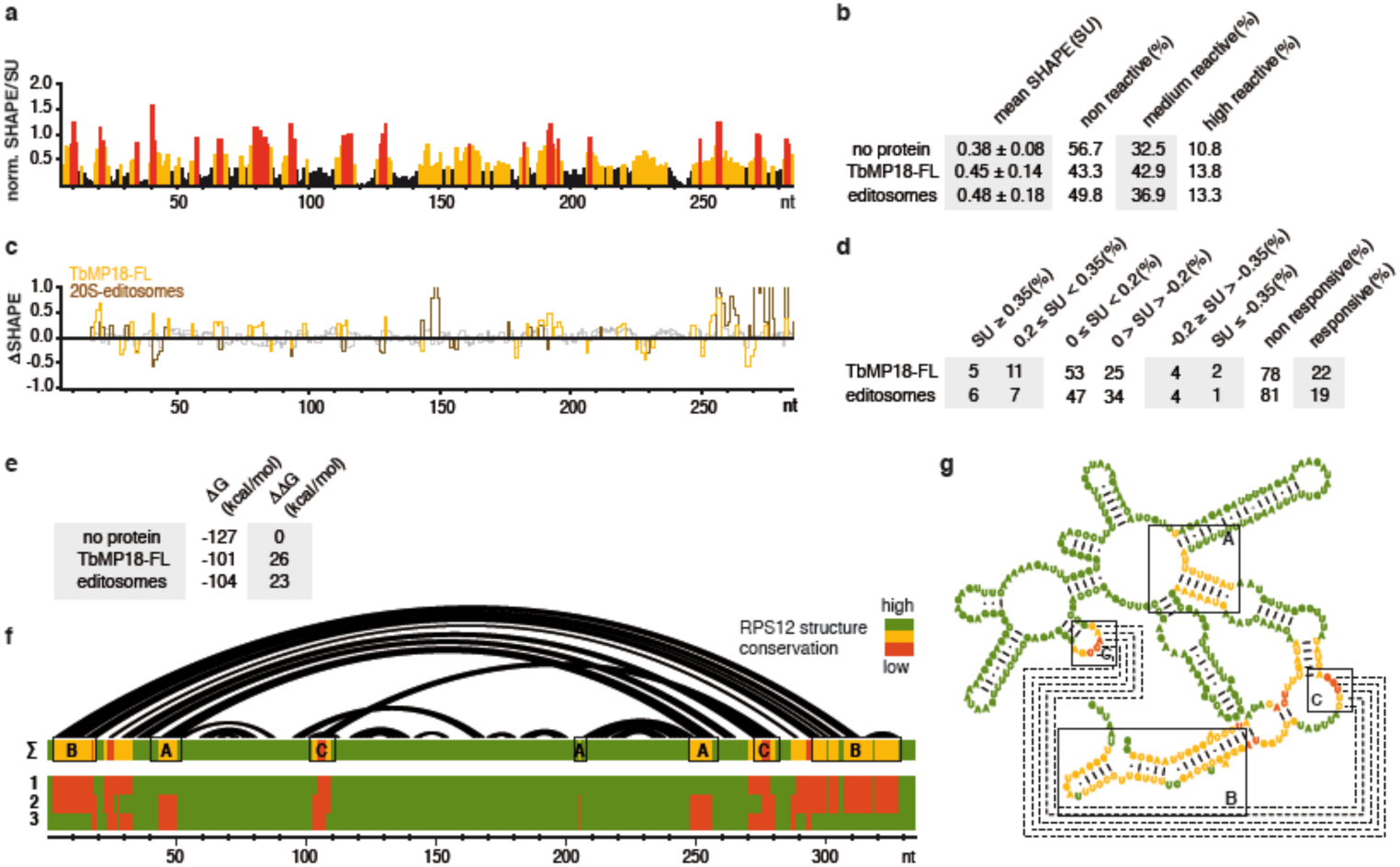
SHAPE-chemical modification of RPS12 pre-mRNA in the presence/absence of TbMP18-FL. (a) Normalized SHAPE-reactivity profile of RPS12 pre-mRNA in the presence of TbMP18-FL. Black: unreactive (SU<0.35), yellow: moderately reactive (0.35≤SU<0.8), red: highly reactive (SU≤0.8) nucleotide positions. (b) Statistical summary of SHAPE-reactivity data. Mean SHAPE-reactivities were calculated from minimally three experiments and are listed ±SD. SU: SHAPE-unit. (c) Difference (Δ)SHAPE-reactivity plots: Yellow: plus/minus TbMP18-FL. Brown: plus/minus 20S editosomes. Non-responsive nt-positions are in grey. (d) Statistical summary of the ΔSHAPE-reactivities. (e) Calculated Gibbs free energies (ΔG) and ΔΔG-values of RPS12 pre-mRNA in the presence/absence of TbMP18-FL and 20S editosomes using the SHAPE-reactivities as pseudo free energy constraints. (f) Arc representation of the basepairing pattern of pre-edited RPS12-RNA in its free conformation. Pairwise comparison of the structure conservation from high (green) to medium (yellow) to low (red). Row 1: free RNA/+editosomes, row 2: free RNA/+TbMP18-FL. Row 3: +editosomes/+TbMP18-FL. ∑: cumulative score to demonstrate the conservation in all three structures. Boxed areas (A, B, C) indicate sequence domains with structural differences. (g) Secondary structure of free RPS12 pre-mRNA with colors indicating the structure conservation as in (F), emphasizing boxed areas A, B, C. Stippled lines: RNA-pseudoknot.

Using the SHAPE-reactivities as pseudo free energy values we calculated minimal free energy (MFE)-structures for the RPS12-RNA in both conformational states (Fig. 5e). As a free RNA, the transcript is characterized by a Gibbs free energy (ΔG) of -127kcal/mol. In the presence of TbMP18-FL the RNA adopts a fold of only -101kcal/mol. This demonstrates a destabilization of the RNA with a ΔΔG of -26kcal/mol. A comparison of all probed structures (free RPS12-RNA, TbMP18-FL-bound RNA and 20S editosome-bound RNA) identified that about 70% of the structural details are shared between the three RNA-folds (Fig. 5f). Differences map to three regions (A, B, C in Fig. 5f,g) involving a pseudoknot and the two termini of the RNA. Together, the SHAPE-chemical probing data corroborate the results of the RNaseH-based gDNA-annealing assay and demonstrate that TbMP18-FL refolds and destabilizes the RPS12-transcript with qualitative and quantitative characteristics similar to 20S editosomes.

### Modeling the RNA-refolding domain(s) of the 20S editosome

Lastly, we asked the question whether the collected data, especially the OB-protein interaction data can be used to derive a structural model of the RNA-refolding domain(s) of the *T. brucei* 20S editosome. For that we performed computational modeling using the program PyRy3D (http://genesilico.pi/pyry3d/) specifically designed to calculate low-resolution models of high molecular mass complexes that contain intrinsic disorder. The program allows the usage of experimentally as well as computationally-derived atomic coordinates together with flexible shapes to include disordered substructures. It has been used to model the structures of several proteins and complexes involved in RNA metabolism including the CCR4-NOT complex^38^. PyRy3D-simulations rely on spatial restraints to implement experimentally-derived interaction data and a scoring function fits the individual components into the contour map of the complex. For that we used the cryo-EM structure of the *T. brucei* 20S editosome^39^. As spatial restraints we implemented all above-described OB-fold interaction data as well as all published binary interactions of editosome components including the recent chemical cross-linking data of the *T. brucei* OB-fold proteins^28-30^. Structure coordinates for the individual proteins were taken from Czerwoniec et al., 2015^17^ (and references therein) and all disordered regions were simulated as coarse-grained flexible shapes. Three hundred independent simulations (300000 steps each) were performed and the resulting 300 models were clustered as outlined in the Supplemental Experimental Procedures. Fig. 6a,b show the medoid structure of the largest cluster (54 models). The different proteins fit tightly into the available volume of the 20S editosome EM-density map with a cross-correlation coefficient of 0.64. A dissection of the structurally well-defined parts of the model from the positions of the intrinsically disordered domains of the different proteins is shown in Fig. 7a,b. This identifies that structured protein domains are preferentially located inside the 0.8MDa complex, while all IDP-domains seem to be preferentially located in peripheral regions of the particle (Fig. 7c). This suggests a general bi-partite domain composition of the editosome: an outer shell of intrinsically disordered proteins, which act as RNA-contact and RNA remodeling elements and an inner core of structurally well-defined proteins, which mediate the catalytic reactions of the complex.

**Figure 6.**
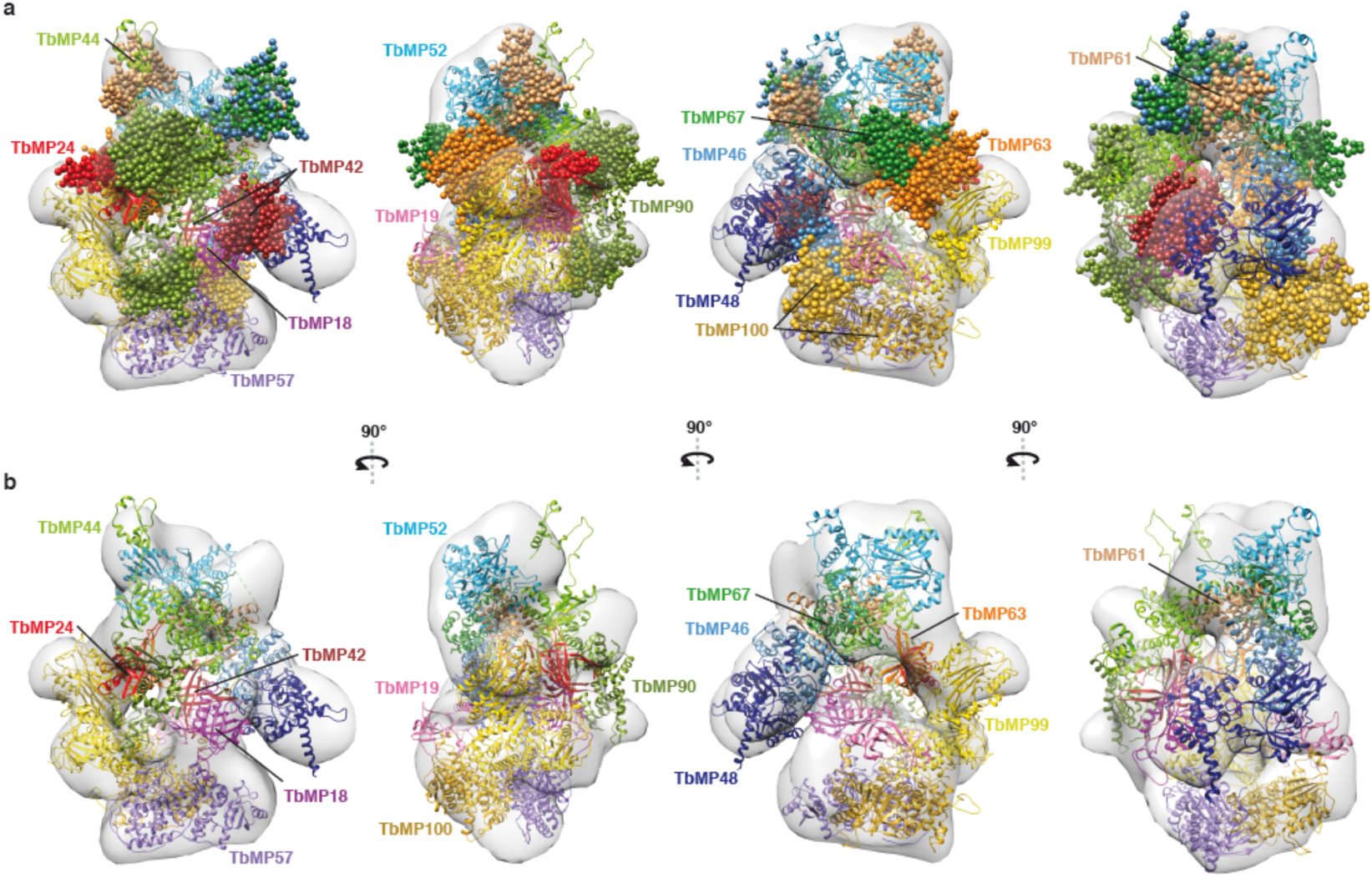
Structure model of the *T. brucei* 20S editosome. (a) Rotational images (90°) of the PyRy3D-derived medoid structure of the 20S editosome. The cryo EM-derived shape of the editosome ^39^ is shown as a gray surface in the background at a contour level of 0.0419. Individual proteins are colour coded and labelled. Intrinsically disordered protein domains are shown as clustered spheres with each sphere representing a Cα-position (diameter 0.19nm). All distance restraints used in the calculation are listed in Supplementary Table S5. PyRy3D can be accessed at (http://genesilico.pi/pyry3d/). The model can be downloaded at. ftp://ftp.genesilico.pl/iamb/models/RNAeditosome3D. (b) 20S editosome model as in (a) omitting the intrinsically disordered protein domains for clarity.

**Figure 7.**
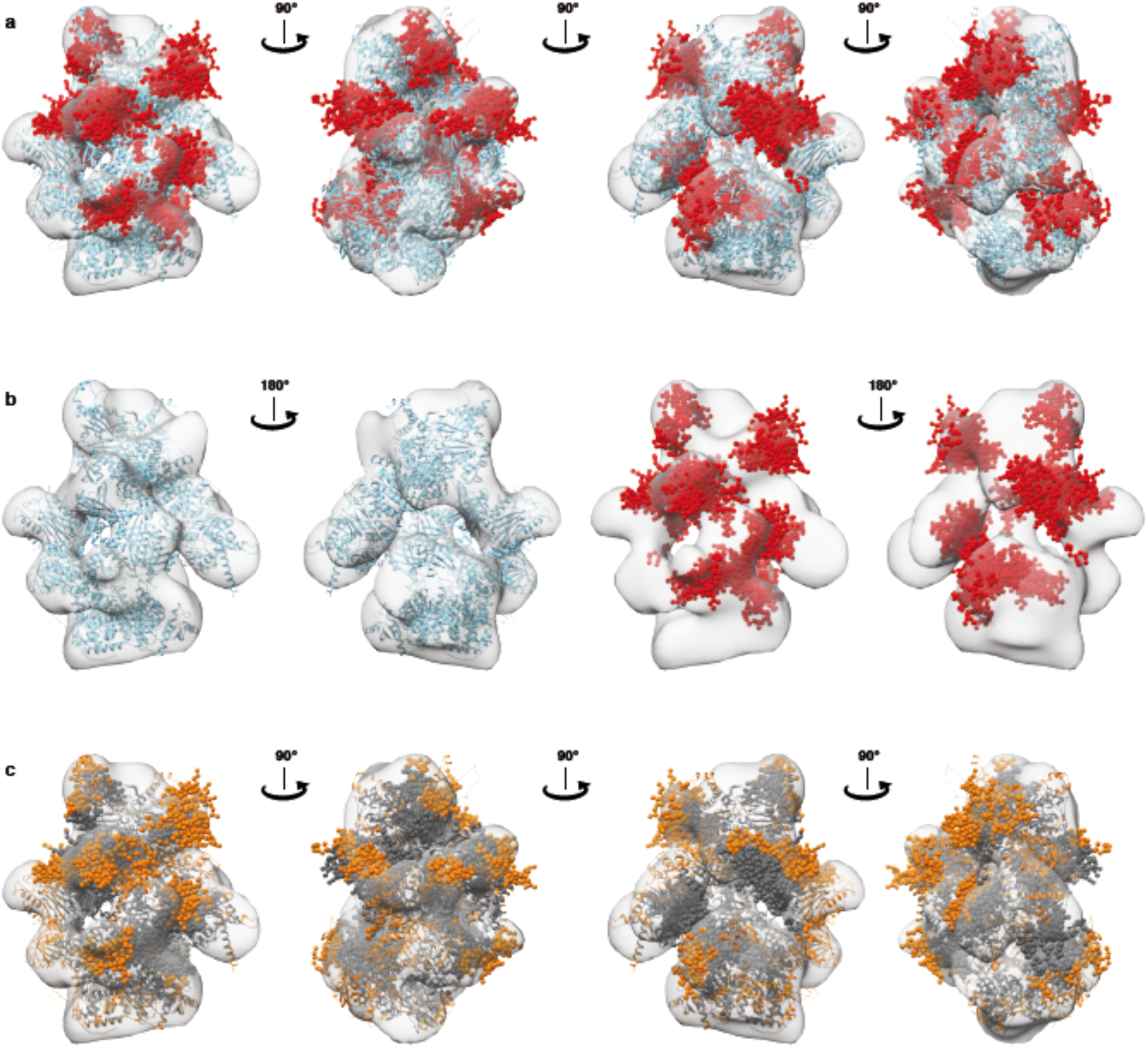
Order/disorder distribution in the 20S editosome. (a) Rotational images (90°) of the PyRy3D-derived 20S editosome model specifically emphasizing the intrinsically disordered regions as clustered spheres in red and structurally ordered regions in light blue. The cryo-EM-based shape of the complex ^39^ is shown as a gray surface (contour level 0.0419). (b) Rotational images (180°) of the editosome model showing only the ordered (light blue) or disordered (red) regions of the complex. Ordered protein domains have a surface/core-ratio of 30/70. IDP-domains are characterized by a value of 40/60. (c) Rotational images (90°) of the editosome model emphasizing surface-located protein domains in orange and structurally “buried” residues in grey.

## DISCUSSION

Editosomes execute a complex-inherent RNA-chaperone activity to remodel the highly folded structures of mitochondrial substrate pre-mRNAs^3,4^. The activity has been characterized to simplify the folding landscape of the different RNAs to facilitate the annealing of gRNAs as templates in the reaction. While several of the protein components of the editosome have been identified to catalyze defined steps of the editing cycle^1^, no editosomal protein has as of yet been recognized to mediate the RNA-chaperone function.

Here we provide evidence that the OB-fold proteins of the editosome execute RNA-remodeling activity. For that we generated recombinant versions of the different proteins and identified that they are highly unstructured. Furthermore, our CD-measurements support computational predictions suggesting that the OB-fold proteins of the *T. brucei* editosome are in large parts intrinsically disordered ^17^. The disordered regions not only locate outside of the OB-fold domains, they can also be found in the loop regions of the different OB-folds and here specifically in the loop sequences between β-sheets β2 and β3 (L_23_) (Fig. 1c). We also confirmed previous observations^28–30^ that the different proteins have a high propensity to homo- and hetero-oligomerize using their OB-folds as interaction domains. Interestingly, all identified pair-wise interactions involve TbMP18, the smallest of the editosomal OB-fold proteins, possibly implicating a key role in the assembly of the editosomal complex.

Importantly, all homo-and hetero-oligomeric complexes showed RNA-chaperone activity. The activities correlate with the surface size of the different proteins indicating that the process is primarily surface-driven with perhaps cumulative characteristics: every OB-fold protein provides a defined value of RNA-unfolding activity, which together generates the overall activity of the editosome. This is supported by the fact that editing-active editosome preparations showed the highest activity. In addition, all FL-constructs displayed higher refolding activities in comparison to the OB-fold-only polypeptides suggesting that the chaperone activity is primarily located within the disordered regions of the proteins. This represents a different situation to the yeast RRP44 protein, where the RNA-unwinding reaction is mediated by a cluster of three OB-folds^31,32^. However, as anticipated, the refolding reaction induces a destabilization of the substrate RNAs, thereby generating RNA-minimal free energy structures of reduced thermodynamic stability. As such, the individual OB-fold proteins execute a refolding reaction that is biochemically and structurally equivalent to the activity of the editosomal complex.

As a consequence of the described results, we propose a scenario in which the OB-folds and the IDP-sequences of the different proteins execute separate but interdependent functions: The OB-folds primarily act as docking modules to assemble an “OB-core” within the editosome as previously discussed^29^, which positions the IDP-sequences of the different proteins on the surface of the catalytic complex where they function to bind and refold mitochondrial pre-mRNAs. Thus, our data add to the functional assignment of the protein inventory of the editosome and within this context it is important to note that about half of the 12 core proteins of the editosome are allocated to remodel RNA. This indicates the importance of the process within the editing reaction cycle. Of course, we can not exclude that IDP-sequences of other, non-OB-fold editosomal proteins such as TbMP90, TbMP67, TbMP61 or TbMP100 might contribute as well^17^.

Intrinsic disorder is a common feature of many RNA-binding proteins^40,41^. It provides the functional advantage of allowing multiple contacts during the initial binding reaction while at the same time enabling sufficient conformational flexibility to target different RNA-ligands^41^. In addition, intrinsic disorder has a kinetic advantage since an increased capture radius results in a higher binding probability. This mechanism is known as “fly-casting mechanism”^42^. It accounts for the fact that structurally flexible protein sequences can enlarge their effective “reach”, which is of special importance for membrane-bound macromolecular complexes especially in crowded solvent conditions. RNA-editing takes place in the highly crowded mitochondria of African trypanosomes and preliminary evidence for an attachment of the editosome to mitochondrial ribosomes and/or the mitochondrial inner-membrane have been discussed^43,44^. We propose that the energy for the RNA-refolding reaction carried out by the editosome follows an “entropy transfer”-mechanism^40^. The binding of a substrate RNA induces a structural re-orientation of the OB-fold proteins, which results in a loss of conformational entropy. This entropy is used to remodel the bound RNA in agreement with the observation that the reaction does not require ATP^3^. Our data further suggest that the pre-edited substrate mRNAs likely interact with the editosome by two successive RNA-binding modes. As in the case of the CBP2/bI5 group I intron interaction^45^ an initial, rather non-specific interaction induces a set of conformational fluctuations in the bound RNA, which is followed by a slow, specific binding mode that stabilizes the editing-competent RNA conformation. Support for such a scenario comes from the observation that flexibility restrictions in model editing substrate RNAs inhibit RNA editing *in vitro*^46^.

As a consequence of our hypothesis we postulate that the IDP-domains of the different OB-fold proteins are located on the surface of the 20S editosome. To provide a structural context for the assumption we performed a “coarse-grained” modeling of the structure of the *T. brucei* editosome using published cryo-EM data^39^ and all OB-fold interaction data as spatial restraints^28-30^. Interestingly, while the model agrees with the anticipated surface location of the different IDP-domains on the editosome, it also suggests a clustering of all structurally well-defined proteins in the center of the complex. Thus, the complex separates an inner core of catalytically active editosome components from an outer shell of ID-proteins that act as RNA-contact and RNA-remodeling elements. Interestingly, a similar separation of structurally defined protein regions from intrinsically disordered domains was also identified for the human spliceosome^47^. Thus, the described organization might reflect a more general structural scenario of macromolecular RNP-machineries.

## METHODS

Full experimental procedures are provided in the Supplemental Experimental Procedures.

### Expression and purification of recombinant OB-fold proteins

DNA-sequences of the six *T. brucei* editosomal OB-fold proteins (TbMP81, TbMP63, TbMP42, TbMP24, TbMP19, TbMP18) were PCR-amplified from *T. brucei* genomic DNA (strain Lister 427)^48^ using the DNA-oligonucleotide primers listed in Supplementary Table S3. Mitochondrial targeting sequences were predicted using MitoProt^49^ and were omitted from the constructs. Proteins were expressed as full length (FL) and OB-fold-only polypeptides and contained a cleavable N-terminal hexa-histidine (His_6_)-tag: TbMP19-FL (aa 18-170), TbMP63-FL (aa 53-587), TbMP81-FL (aa 56-762), TbMP24-OB (aa 126-246), TbMP24-FL (aa 47-246), TbMP42-OB (aa 245-371), TbMP42-FL (aa 22-371), TbMP63-OB (aa 472-587) and TbMP81-OB (aa 626-762). Details of the expression and purification are described in the Supplemental Information.

### RNA-chaperone assay and protein surface area calculation

RNA-chaperone activity assays of the different protein constructs were performed as described by Leeder et al., 2016a^4^ using pre-edited CYb-RNA (1nM) and “gDNA” CYb-5 (100nM) as a pre-mRNA/gDNA-pair. Experiments were conducted using minimally 6 different protein concentrations (10-500nM) in at least two independent experiments to yield ≥ 12 data points. The amount of cleaved RNA was determined by peak-integration and was normalized for each experiment. Editing-active 20S editosomes served as a positive control and were prepared and analyzed as in Leeder et al., 2016a. Data points were fitted to a sigmoidal function: y=L_low_+L_up_/[1+e^(c_1/2_– x/rate)^] (L_low_=lower limit, L_up_=upper limit). The concentration of half maximal RNA cleavage (c_1/2_) represents a proxy for the RNA-chaperone activity. Normalized surface areas were calculated based on the assumption that the proteins have globular shapes. Volume (V) calculations used an average partial specific volume of 0.73cm^3^/g from the molecular masses determined by size exclusion chromatography^50^. Surface areas (A) were calculated as: A=(36πV^2^)^1/3^. All values were normalized to the surface of TbMP81-OB as the smallest protein.

### RNA-synthesis and RNA-structure probing

Pre-edited transcripts of apocytochrome b (CYb) and of ribosomal protein S12 (RPS12) were generated by run off *in vitro* T7-transcription following standard procedures.(^32^P)-labeled RNA preparations were generated by adding α-[^32^P]-UTP to the transcription mix. Selective 2’-hydroxyl acylation analyzed by primer extension (SHAPE) was conducted as in Leeder et al., 2016^4^ using 1-methyl-7-nitroisatoic anhydride (1M7) as the modification reagent. After the RNA-refolding step a 150-fold molar excess (300pmol) of TbMP18-FL was added to the reaction mix and incubated at 27°C for 30min. Raw electrophoretic traces were analyzed using SHAPEfinder^51^ utilizing the boxplot approach to determine the number of statistical outliers. Normalized SHAPE-reactivities were generated by averaging a minimum of 3 independent experiments and were used as pseudo-Gibbs free energies to calculate RNA 2D-structures for a temperature of 37°C using RNAstructure v5.6^52,53^. ShapeKnots^54^ was used to search for pseudoknots using The default parameters p1=0.35 kcal/mol and p2=0.65 kcal/mol.

### Structure modeling of the 20S editosome

Models of the *T. brucei* 20S editosome structure were generated using PyRy3D (http://genesilico.pi/pyry3d/) together with the cryo-EM density map EMDB:1595^39^. Details of the modeling procedure are specified in the Supplementary Information.

## ACKNOWLEDGEMENTS

We thank Matthias Leeder for his help conducting the SHAPE-experiments and Wim Hol (University of Washington, Seattle) for plasmid constructs. This work was supported by the German Research Foundation (DFG-SFB902) and the Dr. Illing Foundation for Molecular Chemistry to H.U.G. The modeling work was supported by the Polish Ministry of Science and Higher Education (grants 0083/IP1/2011/71 and N N301 123138) and by the European Research Council (StG grant RNA+P=123D to J.M.B.). M. D. was supported by the KNOW RNA Research Centre in Poznan (grant 01/KNOW2/2014). J.M.B. was also supported by the ‘Ideas for Poland’ fellowship from the Foundation for Polish Science.

### AUTHOR CONTRIBUTIONS

H.U.G. conceived and supervised the project. C.V. and E.K. conducted the experiments. J.M.B. supervised the computational analysis of the editosome structure. M.D., J.M.K., and A.C. carried out the modeling. P.B. carried out computational analyses of intrinsic disorder. All authors contributed to the analysis and interpretation of the data. H.U.G. and C.V. wrote the manuscript with input from all authors.

### COMPETING FINANCIAL INTERESTS

The authors declare no competing financial interests.

